# Assembly of a functional ribozyme from short oligomers by enhanced non-enzymatic ligation

**DOI:** 10.1101/700229

**Authors:** Lijun Zhou, Derek K. O’Flaherty, Jack W. Szostak

## Abstract

The non-enzymatic replication of the primordial genetic material is thought to have enabled the evolution of the first ribozymes, leading to early forms of RNA-based life. However, the reported rate of chemical RNA ligation is extremely slow. Here we show that the rate of ligation can be greatly enhanced by employing a 3′-amino group at the 3′-end of each oligonucleotide, in combination with an *N*-alkyl imidazole organocatalyst. These modifications allow the rapid copying of long RNA templates by multi-step ligation of tetranucleotides, as well as the assembly of long oligonucleotides from short template splints. Our work shows that a functional RNA ligase ribozyme can be assembled from relatively short oligonucleotides, demonstrating a transition from non-enzymatic ligation to enzymatic ligation. We suggest that the genomes of primitive protocells could have consisted of relatively easily replicated oligonucleotides as short as 10 to 12 nucleotides in length.

## Main Text

The RNA world hypothesis posits an early stage in the evolution of life during which RNA was both the carrier of genetic information and the catalyst of cellular reactions^1–4^. This model is supported by the *in vitro* evolution of ribozymes with RNA polymerase activity^5–7^. However, prior to the emergence of the first RNA replicase ribozyme, genetic polymers must have relied on non-enzymatic self-replication processes to survive^8^. The current lack of an efficient pathway for non-enzymatic RNA replication is the main obstacle towards building a model protocell capable of growth, division, and therefore Darwinian evolution^9^. Recent progress in template copying chemistry with activated mononucleotide substrates has enabled the copying of mixed-sequence RNA templates up to 7 nt long in solution and 5 nt long inside a model protocell^10,11^. In more artificial scenarios, it has been possible to copy bead-immobilized templates up to 12 nt in length, completing a functional ribozyme sequence^12^. In spite of this progress, it is not yet possible to copy, in an effective and prebiotically plausible manner, RNA templates long enough to encode ribozymes that might enable RNA catalyzed self-replication processes^13,14^.

As an alternative to mononucleotide polymerization, templates might be copied by the ligation of short oligomers to generate longer products^15^. However, the slow rate and low yield of template-directed RNA ligation^12,16^ has precluded the copying of long, functional sequences by ligation. In an effort to overcome this problem, the Orgel and von Kiedrowski laboratories used EDC driven ligation of oligonucleotides with 3′-amino-RNA or 3′-phosphoryl-DNA to demonstrate the copying of tetramer and hexamer templates by the ligation of dimer and trimer substrates respectively^17,18^. Unfortunately, no example of long RNA copying with rapid chemical ligation has been reported in the subsequent 30 years. Recently, the Hud laboratory demonstrated the *N*-cyanoimidazole driven ligation of DNA tetramers into long polymers with the help of intercalators to stabilize the tiled tetramer arrays^19^. However this approach failed with RNA tetramers, and no prebiotically plausible intercalator has been reported. The Richert group has recently shown that the rapid cyclization of activated di- and tri-nucleotides significantly limits the ligation of such substrates^20^. Despite these challenges, template copying by ligation remains attractive, as it requires fewer reaction steps to copy a template of a given length. A ligation approach could also overcome other problems that are severe for monomer polymerization, such as copying through secondary structures, copying A/U rich sequences^21^, and copying the last nucleotide of a template^22^. Moreover, short oligomers would have been prebiotically accessible, as inevitable products of reactions between activated monomers both on and off template^23^. In light of these advantages, further studies of non-enzymatic ligation are needed to evaluate the potential of ligation mediated pathways for template assembly and copying.

Here we explore the use of highly reactive oligonucleotide substrates in non-enzymatic template directed ligation. We use short oligoribonucleotides terminated with 3′-amino-2′,3′-dideoxy-ribonucleotides as substrates, because the 3′-amino moiety is a stronger nucleophile than the 3′-hydroxyl of ribonucleotides. Although 3′-amino oligonucleotides are not prebiotically realistic, they are an excellent model for the study of how RNA oligonucleotides might behave if the nucleophilicity of the 3′-hydroxyl could be enhanced, for example by delivery of a catalytic metal ion to the reaction center. We used 2-methylimidazole for 5′-phosphate activation, together with the organocatalyst 1-(2-hydroxyethyl)imidazole (HEI) to provide enhanced activation^24,25^. Together, these modifications allow for the non-enzymatic copying by ligation of templates up to at least 32 nt in length. We also show that oligonucleotides only 4-nt long can assemble into products longer than 140 nt. Finally, we show that a functional RNA ligase ribozyme can be assembled by the splinted ligation of sets of 10-12 nt long oligonucleotides.

We began by quantitatively assessing the effects of a 3′-amino vs. a 3′-hydroxyl nucleophile on the rate of template-directed oligonucleotide ligation. Using an RNA primer containing either a canonical ribonucleotide or a 3′-amino-2′,3′-dideoxyribonucleotide residue at the 3′-end (Fig. 1a, Fig. S1), we measured the rate of primer ligation as a function of concentration of the ligator oligonucleotide, a 2-methylimidazole activated RNA tetramer. The observed maximal rate (*k*_*obs max*_) of the RNA ligation reaction in the presence of 100 mM Mg^2+^ was 0.027 h^−1^ (Fig. 1b, Fig. S2a). RNA ligation in the absence of Mg^2+^ was too slow to measure, while the ligation rate continued to increase up to ~0.1 h^−1^ at 1.6 M Mg^2+^(Fig. S3). Under the same 100 mM Mg^2+^ conditions, substitution of the last nucleotide on the RNA primer with a 3′-amino-2′,3′-dideoxyribonucleotide enhanced the ligation rate by over 50-fold; giving a *k*_*obs max*_ of 1.6 h^−1^ (Fig. 1c, Fig. S2b). Notably, without Mg^2+^, *k*_*obs max*_ reached an even higher value of 5.1 h^−1^ (Fig. 1c, Fig. S2c). Our results show that Mg^2+^ accelerates the N3′–P5′ ligation rate only at low concentrations of the ligator oligonucleotide. When the concentration of the ligator is sub-saturating, Mg^2+^ stabilizes duplex formation and improves the *k*_*obs*_. At higher, saturating concentrations of tetramer or with a longer ligator, such that the template is saturated even in the absence of Mg^2+^, the presence of Mg^2+^ lowers *k*_*obs max*_, for reasons that remain unclear (Fig. S3-S7).

**Figure 1:**
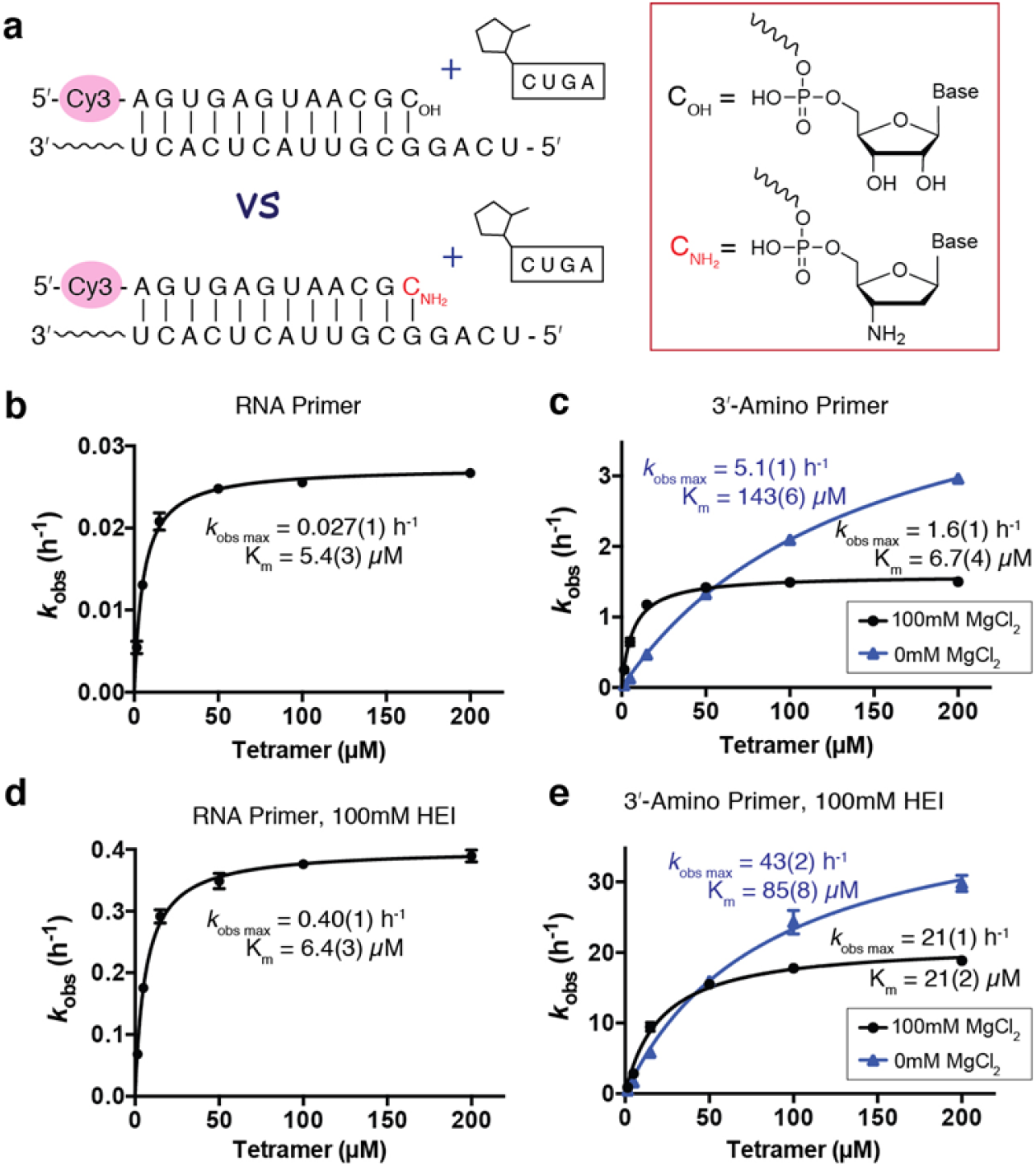
Kinetic studies of non-enzymatic ligation reactions. All experiments were carried out with 2 μM primer, 4 μM template, 200 mM HEPES, pH 8.0, and 100 mM MgCl_2_ unless otherwise noted, and with the indicated concentration of the tetramer ligator. (a) Illustration of experimental design. The 5′-fluorescently labeled primer contains either a ribonucleotide (C_OH_) or a 3′-amino-2′,3′-dideoxyribonucleotide (C_NH2_) at the 3′-terminus. The template is complementary to the primer with a 5′-UCAG overhang. The ligator is 2-MeImp-CUGA. (b)-(e) show the ligator concentration dependence of the ligation rate under different conditions: (b) RNA primer; (c) 3′-amino primer; (d) RNA primer with 100 mM HEI; (e) 3′-amino primer with 100 mM HEI. The black circles: reactions with 100 mM MgCl_2_. Blue triangles: reactions with no MgCl_2_. Data points are reported as the mean +/− s.d., n ≥3.

The organocatalyst 1-(2-hydroxyethyl)imidazole (HEI) enhances the non-enzymatic polymerization rate of imidazole activated monomers^24,26^. HEI is thought to act as a nucleophilic catalyst that exchanges with the 2-methylimidazole moiety on the activated nucleotides, generating a more reactive imidazolium leaving group. As expected, the presence of HEI increased the rate of RNA ligation in the presence of 100 mM Mg^2+^ by more than ten-fold, such that *k*_*obs max*_ reached 0.40 h^−1^ (Fig.1d, Fig. S8a). Similarly, HEI also increased the rate of the N3′–P5′ ligation by more than ten-fold to a *k*_*obs max*_ of 21 h^−1^ and 43 h^−1^, in the presence and absence of 100 mM Mg^2+^ respectively (Fig. 1e, Fig. S8b, Fig. S8c).

Encouraged by the greatly enhanced ligation rate resulting from the combination of a 3′-amino nucleophile and the HEI organocatalyst, we applied these advances to the copying of longer RNA templates by multi-step ligation. We first tested the copying of a 32 nt RNA template containing eight 5′-UCAG-3′ repeats (Fig. 2a) in the presence of 400 μM of the tetramer 2-MeImp-CUGA-3′-NH_2_. With HEI, we observed a ~20% yield of full-length product within 10 minutes and more than 60% within 40 minutes. Consistent with our previous results, Mg^2+^ inhibited ligation, leading to increased levels of a ladder of stalled products corresponding to the stepwise addition of tetramers. After two hours, we did not observe any further increase in the yield of full-length product, and the stalled products remained unchanged in intensity. To test the hypothesis that the stalled products arise from templates occupied by hydrolyzed (i.e. unactivated) downstream ligator oligonucleotides, we repeated the ligation experiments but added multiple aliquots of the coupling reagent EDC after the initial stage of the reaction was largely complete. In the absence of Mg^2+^, this strongly decreased the background of stalled products and significantly increased the yield of full-length product (Fig. S9). The effect of EDC was much smaller in the presence of Mg^2+^, possibly due to Mg^2+^-enhanced EDC hydrolysis. We suggest that more prebiotically plausible chemistry that could re-activate hydrolyzed ligator oligonucleotides, such as the recently described isonitrile activation pathway^27^, might also significantly increase the yield of full-length product in multi-step template copying reactions.

**Figure 2:**
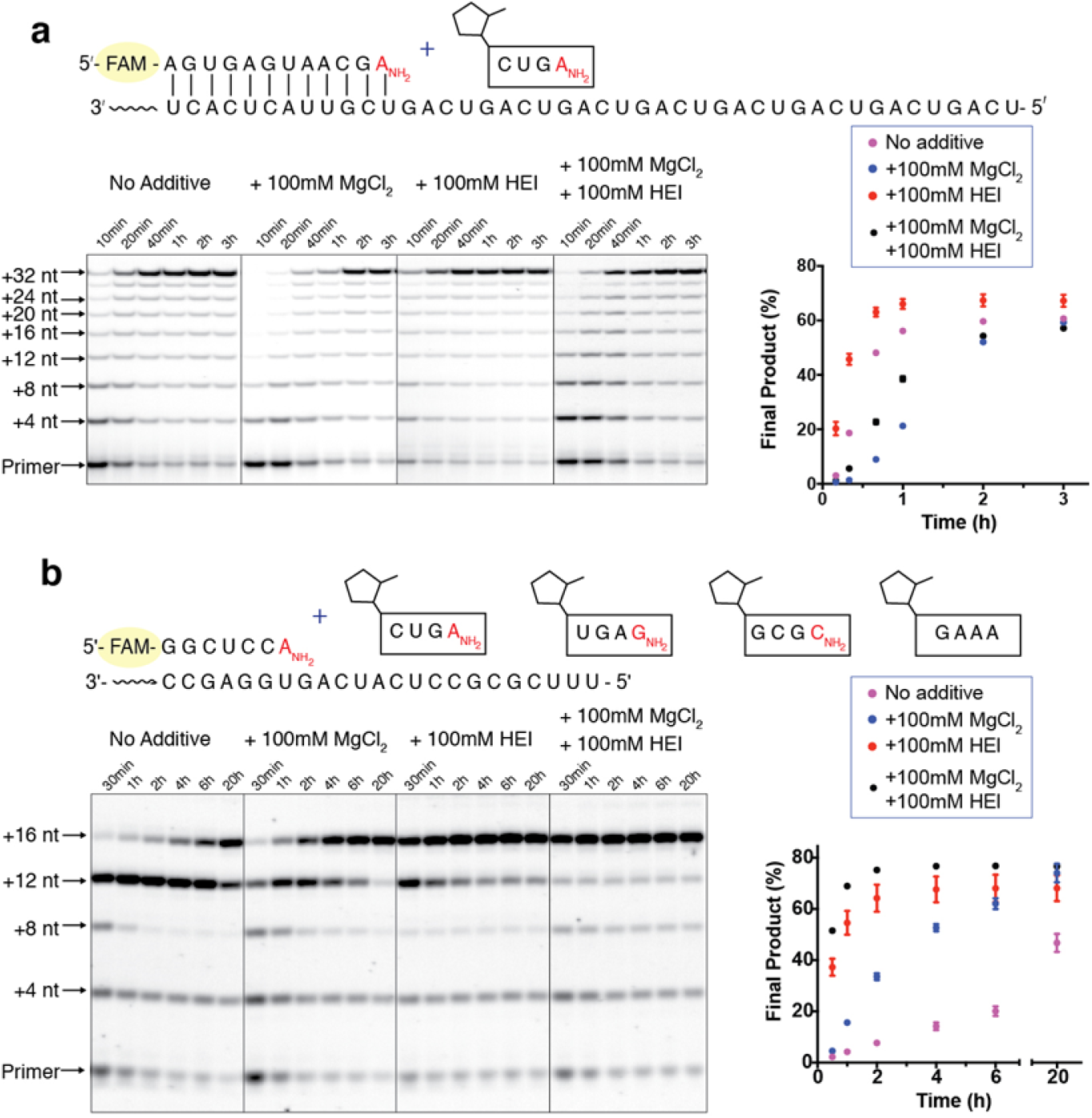
Non-enzymatic oligonucleotide ligation *via* N3′–P5′ linkage formation can rapidly copy a long RNA template. (a) copying of a 32 nt long repeating sequence by the tetramer 2-MeImp-CUGA_NH2_. Reactions contained 1.5 μM primer, 3 μM template, 200 mM HEPES, pH 8.0, 400 μM 2-MeImp-CUGA_NH2_. (b) copying of a 16 nt long sequence by ligation of four different tetranucleotides. Reactions contained 2 μM primer, 3 μM template, 200 mM HEPES, pH 8.0, 50 μM 2-MeImp-CUGA_NH2_, 50 μM 2-MeImp-UGAG_NH2_, 50 μM 2-MeImp-GCGC_NH2_, 50 μM 2-MeImp-GAAA. Data points are reported as the mean +/− s.d., n≥3.

We then examined the copying of a 16 nt template with a non-repeating sequence, in the presence of four different tetramers at 50 μM each (Fig. 2b). As before, full-length product was formed rapidly in the presence of HEI, with ~35% full-length product observed by 30 minutes and more than 60% at 2 hours. In this system, Mg^2+^ significantly improved the rate of full-length product formation. This effect was particularly notable for the addition of the last tetramer 2-MeImp-GAAA, where Mg^2+^ likely facilitated binding of this A-rich tetramer. Thus, the combination of a 3′-amino nucleophile, HEI catalysis and Mg^2+^ allows for the rapid and high yielding copying of mixed sequence RNA templates.

Although multi-step ligation allows the copying of long RNA templates, the thermal stability of the resulting long RNA duplexes makes it difficult to separate the strands so that the product strand could either fold into an active ribozyme or act as a template for the next round of copying^9^. We therefore asked if multi-step ligation could be directed by short RNA template splints which could dissociate more easily. Enzyme-catalyzed splinted ligation^28^ is widely used to generate RNA products that are much longer than the starting materials, hinting at a plausible chemical pathway for emergence of the first long ribozyme sequences or templates in the pre-RNA world. We therefore asked whether long oligonucleotides could assemble starting from the splint template 5′-AGUC-3′, and the activated tetramer 2-MeImp-CUGA-3′-NH_2_ (Fig. 3a). The template and the ligator can assemble into tiled arrays held together by two base pairs on each side of each tetramer. To monitor the non-enzymatic ligation reaction, we used a 5′-fluorescently-labeled primer containing a 3′-amino terminus hybridized to a template with a 5′-AG overhang. We tested the reaction with and without 100 mM MgCl_2_ and 100 mM HEI. In this experiment, the limiting factor is the weak binding between the two-nucleotide overlaps. We therefore conducted the experiment on ice, with 200 mM NaCl, to promote pairing. MgCl_2_ helped to stabilize the tiled arrays and therefore greatly enhanced the long oligo formation, resulting in >140 nt products within 1 day. When using both MgCl_2_ and HEI, >100 nt products formed within 6 hours (Fig. 3b). This experiment serves as a proof-of-concept that non-enzymatic ligation is capable of generating long oligomers without pre-existing long templates.

**Figure 3:**
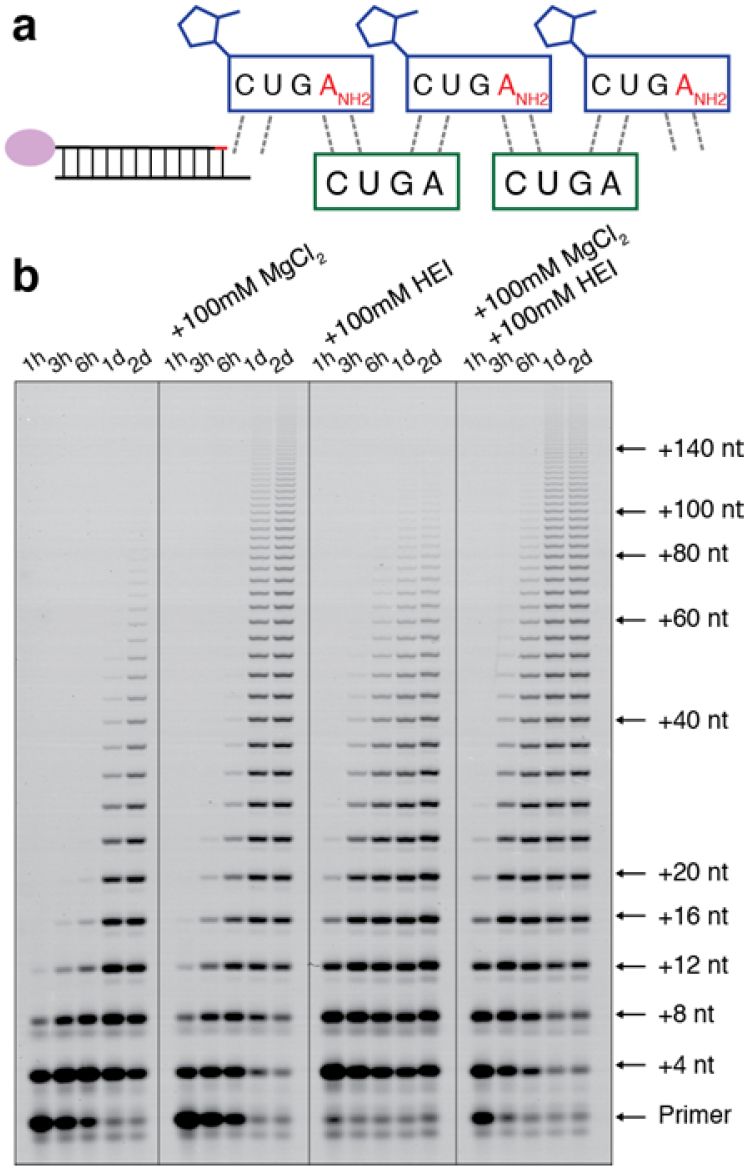
The non-enzymatic ligation reaction of tetramers leading to the formation of long oligomers without a pre-existed long template. (a) Schematic representation of the reaction. The splint template is the non-activated tetramer 5′-AGUC. The ligator is 2-MeImp-CUGA_NH2_. The duplex of 5′-Cy3-labeled primer with 3′-A_NH2_ and 5′-AG-overhang RNA template was used to monitor the reaction. (b) PAGE shown for the ligation reactions leading to the formation of >140 nt products within 1 day. The reactions contained 100 mM HEPES, pH 8.0, 500 μM AGUC, 500 μM 2-MeImp-CUGA_NH2_, 200 mM NaCl, with/without 100 mM MgCl_2_ or 100 mM HEI as noted. The experiments were carried out on ice in triplicate.

Having established that long RNA strands could assemble from short oligonucleotides via splinted ligation, we asked whether non-enzymatic ligation could lead to the formation of functional ribozymes. The nonenzymatic assembly of an RNA ligase ribozyme is a tantalizing prospect, since this type of ribozyme could represent a key intermediate in the transition from the pre-RNA world to the RNA world. Recently, the Holliger group reported the assembly of an RNA polymerase ribozyme by the nonenzymatic ligation of six 20-nt and one 30-nt long 2-methylimidazole activated RNA oligonucleotides in 0.5% overall yield^29^. However, T7 RNA polymerase transcription of RT-PCR amplification product was needed to demonstrate ribozyme activity, possibly because of the failure of the 20-30 nt splints to dissociate from the assembled product. In order to study a simpler and more prebiotically relevant model system, we decided to examine the assembly of a much smaller ligase ribozyme from shorter oligonucleotides. The Joyce group has reported a highly-efficient RNA ligase ribozyme of 52-nt in length, which can ligate a 14-nt 5′-triphosphate RNA substrate to its 3′-end ^30^. We designed an experiment to generate this particular ligase from five different oligomers, varying from ten to twelve nucleotides in length (Fig. 4), with ligation directed by four different 10-nt splint oligonucleotides. Preliminary experiments with DNA splints showed efficient assembly of full-length ribozyme, which upon dilution into ribozyme buffer exhibited the expected ligase catalytic activity at the optimal temperature of 48°C (Fig. S10). Subsequent experiments with RNA splints of the same length and sequence also exhibited excellent assembly of the full-length ligase ribozyme, but upon dilution into ribozyme buffer, no ligase activity was detected at 48°C, presumably due to inhibition by the tightly bound RNA splint oligonucleotides (Fig. S11). Shorter 8-nt splints resulted in a severe loss of ribozyme assembly efficiency. We therefore introduced a number of G:U wobble pairs between the splint-templates and the ligator oligonucleotides, to lower the melting temperature while maintaining specific binding. In addition, one of the four splints was shortened to 9-nt, instead of 10-nt. In this non-enzymatic ligation experiment, the mixture of the splints and ligators gave rise to the 52-nt product at ~50% yield within 6 hrs, and ~66% in 48 hrs (Fig. 5c, Fig. S12). To test the ligase activity of assembled product, we diluted the ligation reaction into ribozyme ligase buffer containing the 5′-triphosphate substrate, and incubated the mixture at 48°C. Strikingly, the 52-nt product of the non-enzymatic ligation reaction could efficiently perform the enzymatic ligation reaction with the 5′-triphosphate RNA substrate (Fig. 5c).

**Figure 4:**
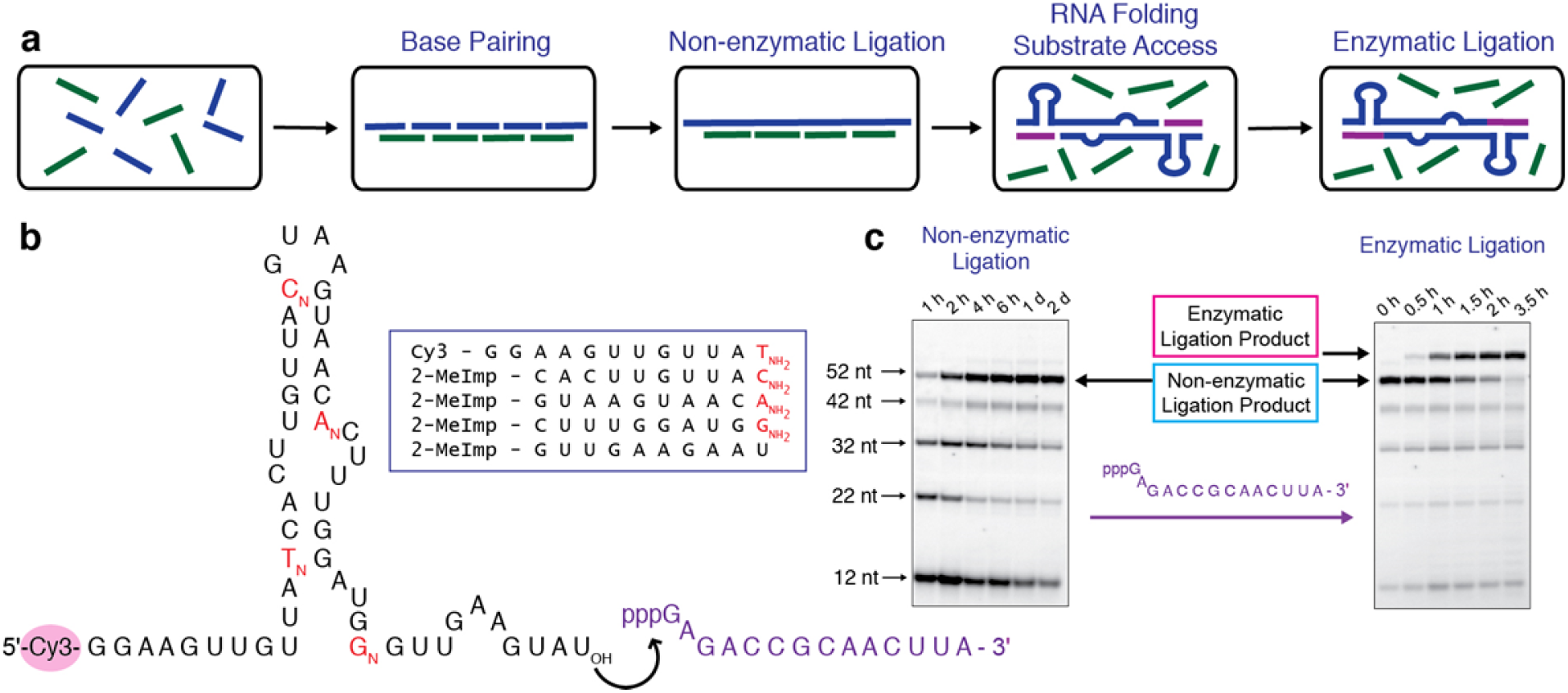
The transition from non-enzymatic ligation to enzymatic ligation. (a) Schematic representation of the experiment design. Oligonucleotides shown in blue take part in multi-step ligation reactions templated by the green splint oligonucleotides to generate the full ligase sequence. Following dilution, the splint oligonucleotides dissociate in the 48 °C ribozyme reaction mixture, allowing folding of the ligase ribozyme. The functional ligase then ligates itself to the 5′-triphosphate oligonucleotide substrate. (b) Proposed secondary structure of ribozyme ligase. Black letters represent ribonucleotides; red letters represent the 3′-amino-2′,3′-dideoxyribonucleotides used in this study. The set of ligator oligonucleotides is shown inside the box. The purple sequence is the 5′-triphosphate substrate of the ribozyme ligase. (c) Left PAGE shows the formation of RNA ligase ribozyme *via* non-enzymatic ligation. The reaction contains 20 μM primer, 25 μM each 2-methylimidazole activated oligonucleotide as shown in (b), 25 μM each of the four different RNA template splints (5′-AAGUGAUAAU-3′, 5′-CUUAUGUAAU-3′, 5′-UAAAGUGUUA-3′ and 5′-CAACCUAUU-3′), 100 mM HEPES, pH 8.0, 100 mM NaCl, 50 mM MgCl_2_ and 100 mM HEI. The ligation reaction was carried out on ice. Right PAGE shows the activity of the product ligase. The non-enzymatic ligation product was diluted 10-fold into the ribozyme reaction buffer: 50 mM bis-tris propane, pH 8.5, 25 mM MgCl_2_ and 3 μM triphosphate substrate. The ligase reaction was performed at 48 °C.

Our work suggests a scenario for the origin of life in which the replicating genomic oligonucleotides in the first protocells were much smaller than previously considered. Because reasonably active ribozyme nucleases are generally larger than 25-50 nucleotides in length, with ligases and polymerases being substantially longer, it had generally been assumed that replicating genomic fragments had to be at least 25 nucleotides in length if not longer. Even when ribozymes are assembled from multiple oligonucleotides, the fragments lengths are typically ≥ 25 nucleotides long. Copying sequences of this length by primer extension remains an unsolved problem, and even if that problem can be solved, complete cycles of replication will require a means of overcoming the thermodynamic stability of such long RNA duplexes so that the separate strands can be copied. In contrast, the full replication of oligonucleotides in the range of 10-12 nucleotides long seems much more plausible, both in terms of template copying and because of the lower melting temperature of such short duplexes. Oligonucleotides only 10-12 nucleotides long are unlikely to exhibit useful catalytic activities individually, but could assemble by multiple splinted ligation reactions into much longer oligonucleotides that could indeed act as efficient ribozymes, as shown by our assembly of an active ribozyme ligase by a series of nonenzymatic ligation reactions.

Demonstrating cycles of replication of genomic RNA oligonucleotides and the concurrent assembly of longer functional ribozymes will require overcoming several obstacles. First, the efficient copying of RNA templates of 10-12 nucleotides length must be achieved either by monomer polymerization, by the ligation of small oligomers, or by some combination thereof. Second, efficient splinted-ligation must be achieved with RNA so that longer functional (but non-replicating) sequences can be assembled. Third, the duplex products of oligonucleotide copying must dissociate in order to allow for iterated cycles of replication. Finally, the affinity of splint-templates for their ligator oligonucleotides must be high enough to allow for efficient ligation, but the affinity of the splints for the ligated product strand must be low enough to allow for splint dissociation, so that the assembled product strand can fold into an active ribozyme. The first two problems would benefit from an effective way of increasing the nucleophilicity of the RNA 3′-hydroxyl, perhaps by some means of delivering an appropriate catalytic metal ion to the reaction center. The third problem might be overcome if either strand displacement or strand separation could be triggered by transient environmental changes, for instance, temperature fluctuations or salt concentration fluctuations following wet-dry cycles. The final problem of achieving splint specificity (so as to correctly program ribozyme assembly) with only moderate affinity was an unexpectedly difficult hurdle in our efforts to demonstrate the assembly of an RNA ligase through nonenzymatic ligation. RNA splints that had only 3-4 nucleotides of complementarity to two ligator oligonucleotides led to slow and/or inaccurate ligation, and thus inefficient assembly. On the other hand, splints with 10-12 nucleotides of complementarity to the ligated product strand failed to dissociate, leading to inhibition of the assembled ribozyme. We found that the introduction of wobble mis-matches allowed RNA splints to accurately direct ribozyme assembly through iterated ligation events without inhibiting the assembled ribozyme. Similarly, DNA splints exhibited excellent specificity with low enough affinity to avoid ribozyme inhibition. Prebiotically, it is possible that the admixture of low levels of deoxy-, arabino- or threo-nucleotides into RNA oligonucleotides could have had the same effect, as well as potentially decreasing duplex stability enough to facilitate multiple cycles of replication. The identification of roles for weakly binding oligonucleotides is reminiscent of our previously described mechanism for the homeostatic modulation of ribozyme activity by short oligonucleotides that inhibit ribozyme folding at high concentrations, but dissociate and thereby lead to ribozyme activation following the dilution that occurs during protocell growth^31^. Solving the related problems of oligonucleotide replication and splint-directed ribozyme assembly may lead to further insights into the origins of life in an RNA world.

## Acknowledgments

J.W.S. is an Investigator of the Howard Hughes Medical Institute. This work was supported in part by a grant (290363) from the Simons Foundation to J.W.S. and by a grant from the NSF (CHE-1607034) to J.W.S. D.K.O. is a recipient of a Postdoctoral Research Scholarship (B3) from the Fonds de Recherche du Quebec−Nature et Technologies (FRQNT), Quebec, Canada, and a Postdoctoral Fellowship from Canadian Institutes of Health Research (CIHR) from Canada. The authors thank Constantin Giurgiu, Dr. Thomas Wright, Seohyun Chris Kim, Dr. Li Li, and Dr. Victor Lelyveld for helpful discussion and technical assistance. The authors thank the Szostak group for helpful feedback.

## Author contributions

All authors contributed to the design of the experiments and to writing the paper. Experiments were conducted by D.K.O and L.Z.

## Data availability statement

All data supporting the findings of this study are available within the Article and its Supplementary Information, or from the corresponding author upon reasonable request.

