## Supplementary Materials for "Assembly of a functional ribozyme from short oligomers by enhanced non-enzymatic ligation"

**This PDF file includes:**

Materials and Methods
Figs. S1 to S12

### Materials and Methods

Chemical synthesis of phosphoramidites

Protected 3′-amino phosphoramidites were prepared according to previously reported procedures[1]. 3′-Amino-2′,3′-dideoxycytidine was purchased from Carbosynth (Campton, UK). 3′-Amino-2′,3′-dideoxyguanosine, 3′-amino-2′,3′-dideoxyadenosine and 3′-amino-2′,3′-dideoxythymidine were purchased from Fisher/Alfa Aesar (Haverhill, MA). Other chemical reagents were purchased from Sigma-Aldrich.

Solid phase synthesis of oligonucleotides

Oligonucleotides were prepared by solid-phase synthesis on an Expedite 8909 or MerMade 6 DNA/RNA synthesizer (Bioautomation, Plano, TX). Chemical reagents were purchased from Glen Research (Sterling, VA) and Chemgenes (Wilmington, MA). RNAs were deprotected by standard methods. Non-dye labeled oligonucleotides were purified by GlenPak columns. Dye-labeled oligos were purified by polyacrylamide gel electrophoresis and desalted by Sep-Pak C18 cartridge from Waters (Milford, MA).

Activated Oligonucleotides

5′-phosphorylated oligonucleotides were activated with 2-methylimidazole as previously described[2,3]. Activated tetranucleotides were purified on Agilent ZORBAX PrepHT columns (Eclipse XDB-C18, 250 × 21.2 mm, 7 µm particle size, P.N. 977250-102), at a flow rate of 15 ml/min, using gradient elution between (A) aqueous 20 mM triethylammonium bicarbonate, pH 7.5, and (B) acetonitrile. Activated tetramers were separated from unactivated tetramers with a gradient of 6% to 11 % B over 14 minutes. Activated decanucleotides were purified on Agilent ZORBAX analytical columns (Eclipse Eclipse Plus C18, 250 × 4.6mm, 5 µm particle size, P.N. 959990-902), at a flow rate of 1 ml/min, using gradient elution between buffers A and B as above. Activated decamers were separated from unactivated decamers with a gradient of 6% to 10 % B over 17 minutes.

High-resolution mass spectrometry

Oligonucleotides were analyzed on an Agilent 1200 HPLC coupled to an Agilent 6230 TOF mass spectrometer. Samples were resolved by IP-RP-HPLC on a 100 mm × 1 mm Xbridge C18 column with 3.5 µm particle size (Waters, Milford, MA) using gradient elution between (A) aqueous 200 mM 1,1,1,3,3,3-hexafluoro-2- propanol with 1.25 mM triethylamine, pH 7.0, and (B) methanol. Samples were analyzed in negative mode from 239 m/z to 3200 m/z with a scan rate of 1 spectrum/s.

Non-enzymatic Ligation Reaction

Unless otherwise indicated, non-enzymatic primer extensions were performed according to previously reported[4] procedures except that we used ligator oligonucleotides instead of mononucleotides. At each time point, 0.5 µl reaction was added to 25 µl quenching buffer, containing 10 mM EDTA, 5-10 µM complementary strand RNA, 0.5 x TBE buffer and 90 % (v/v) formamide. Quenched samples were heated to 95 °C for 1 min and cooled quickly to room temperature (typically 25 °C). Primer extension products were resolved by 20 % (19:1) denaturing PAGE with 7 M urea. For $MgCl_2$ titration experiments, the quenching buffer contained 50 mM EDTA instead of 10 mM. For the experiments in Fig. S6 and Fig. S7, the quenching buffer was pre-heated to 100 °C, to ensure immediate quenching, because at room temperature, the N3′–P5′ ligation of the decamer ligator continues in formamide or 8M urea.

Ligase ribozyme activity test reaction

Non-enzymatic ligation reaction products were stored at -80°C prior to testing the ligase enzymatic activity. Non-enzymatic product samples were thawed, diluted 10-fold in a reaction buffer containing 50 mM bis-tris propane, pH 8.5, 3 μM triphosphate RNA substrate, 25 mM $MgCl_2$ and incubated at 48°C. At each time point, 1 μl reaction was added to 15 μl quenching buffer containing 10 mM EDTA, 0.5 x TBE buffer and 90% (v/v) formamide. The enzymatic ligation products were resolved by 20% (19:1) denaturing PAGE with 7 M urea.

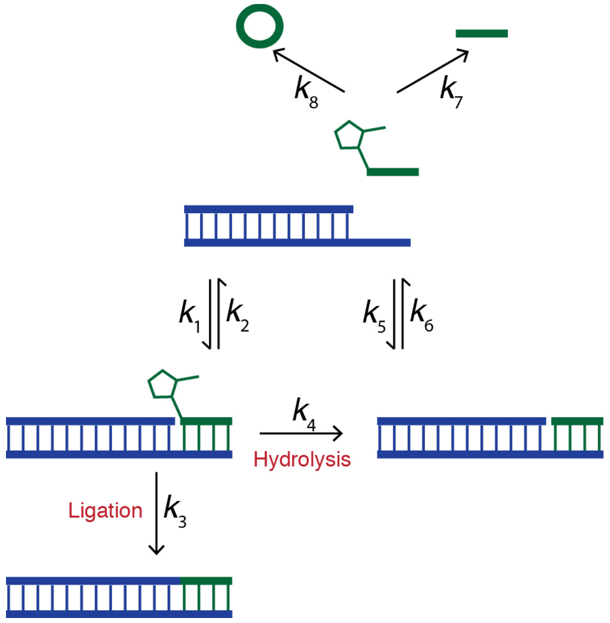


**Fig. S1:**

Kinetic scheme for reactions involved in non-enzymatic oligonucleotide ligation. Hydrolysis reactions, both on and off template, compete with the ligation reaction. Reversible binding of the ligators to the template is followed by competition between ligation and hydrolysis. The fraction of the primer/template complex that ultimately becomes ligated depends upon the dissociation of inactive (hydrolyzed) ligators to allow new incoming activated ligators to bind the template.

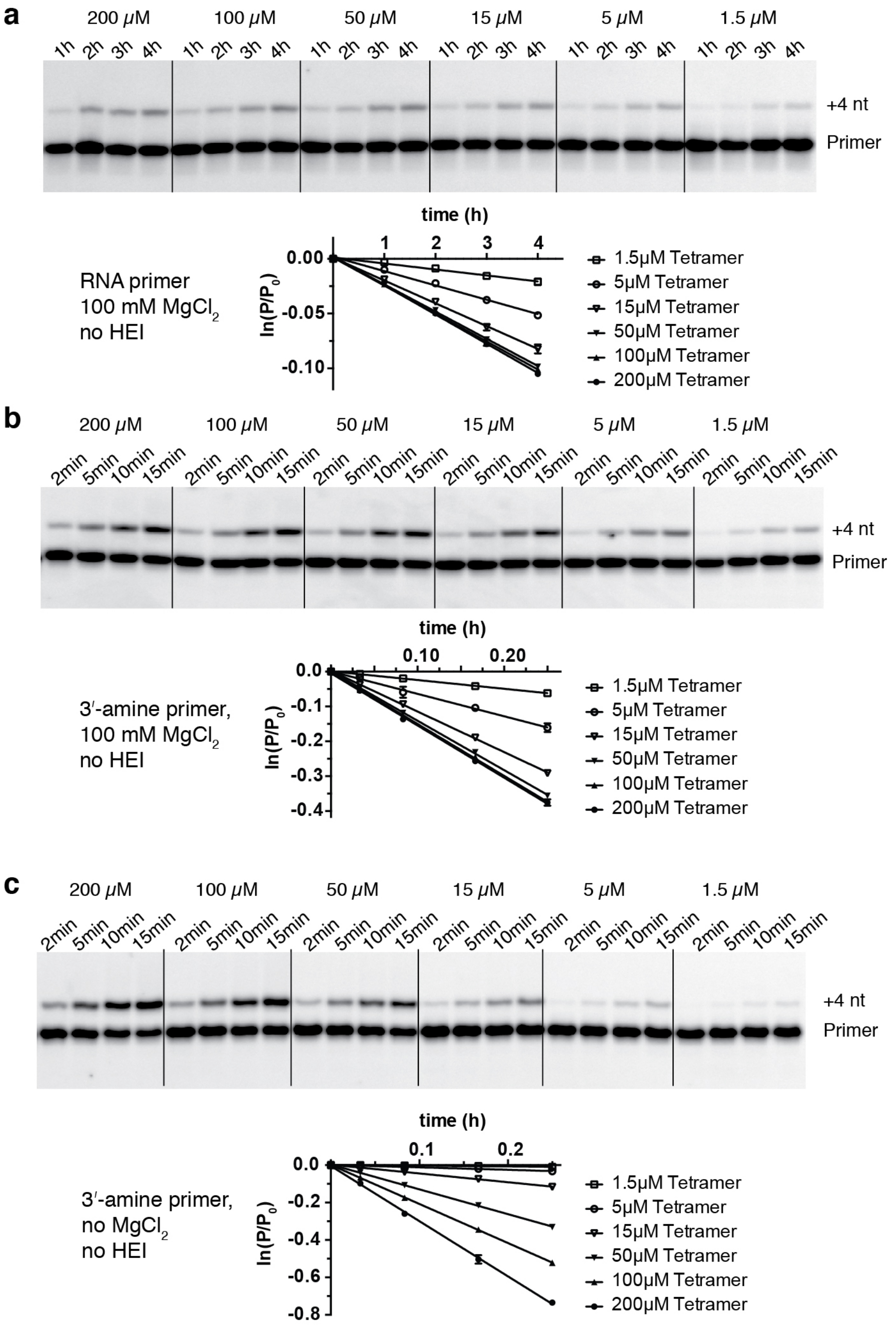


**Fig. S2**

Representative PAGE data and plots of ln($P/P_0$) vs. time for the ligation kinetics described in Figures 1b and1c. (a) Ligation of the all RNA primer to the activated tetramer at the indicated concentrations, in the presence of 100 mM $MgCl_2$, in the absence of HEI. (b) Ligation of the 3′-amino terminated primer to the activated tetramer at the indicated concentrations, in the presence of 100 mM $MgCl_2$, in the absence of HEI. (c) Ligation of the 3′-amino terminated primer to the activated tetramer at the indicated concentrations, in the absence of both $MgCl_2$ and HEI.

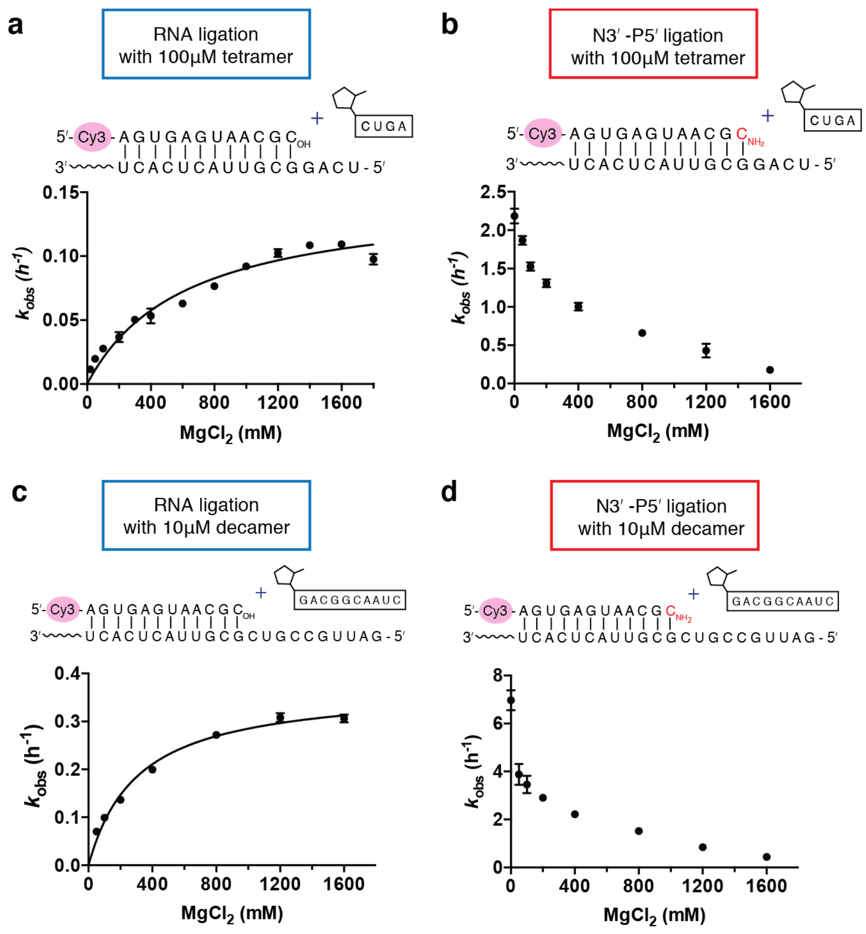


**Fig. S3**

Rates of oligonucleotide ligation reactions as a function of $MgCl_2$ concentration. (a) Ligation of RNA primer with 100 µM tetramer (2-MeImp-CUGA). (b) Ligation of 3′-amino terminated primer with 100 µM tetramer (2-MeImp-CUGA). (c) Ligation of RNA primer with 10 µM decamer (2-MeImp-GACGGCAAUC). (d) Ligation of 3′-amino terminated primer with 10 µM decamer (2-MeImp-GACGGCAAUC). All experiments were carried out with 2 µM primer, 4 µM template, 200 mM HEPES, pH 8.0, 100 µM tetramer or 10 µM decamer, and $MgCl_2$ at the indicated concentration. Data are reported as the mean +/- s.d. from triplicate experiments.

To minimize the effect of $Mg^{2+}$ on the binding of the ligators, we performed reactions with 100 µM tetramer ligator or 10 µM decamer ligator. $Mg^{2+}$ is thought to assist RNA ligation by deprotonating the 3′-hydroxyl nucleophile. As a result, $k_{obs}$ increases with increasing $Mg^{2+}$ concentration, up to 1.4M and 1.2M for the tetramer and decamer ligation, respectively. In contrast, $k_{obs}$ of N3′–P5′ ligation decreases as the concentration of $Mg^{2+}$ increases. We speculate that $Mg^{2+}$ interacts unfavorably with 3′-amine of the primer or the 2-methylimidazole moiety of the ligator. Additionally, $Mg^{2+}$ may affect the conformation of the duplex and disrupt the reaction geometry necessary for N3′–P5′ ligation. Here, we conclude that $Mg^{2+}$ is beneficial for N3′–P5′ ligation only when the concentration of the ligator is too low to afford sufficient binding to the template. Otherwise, the presence of $Mg^{2+}$ decreases the rate of the reaction.

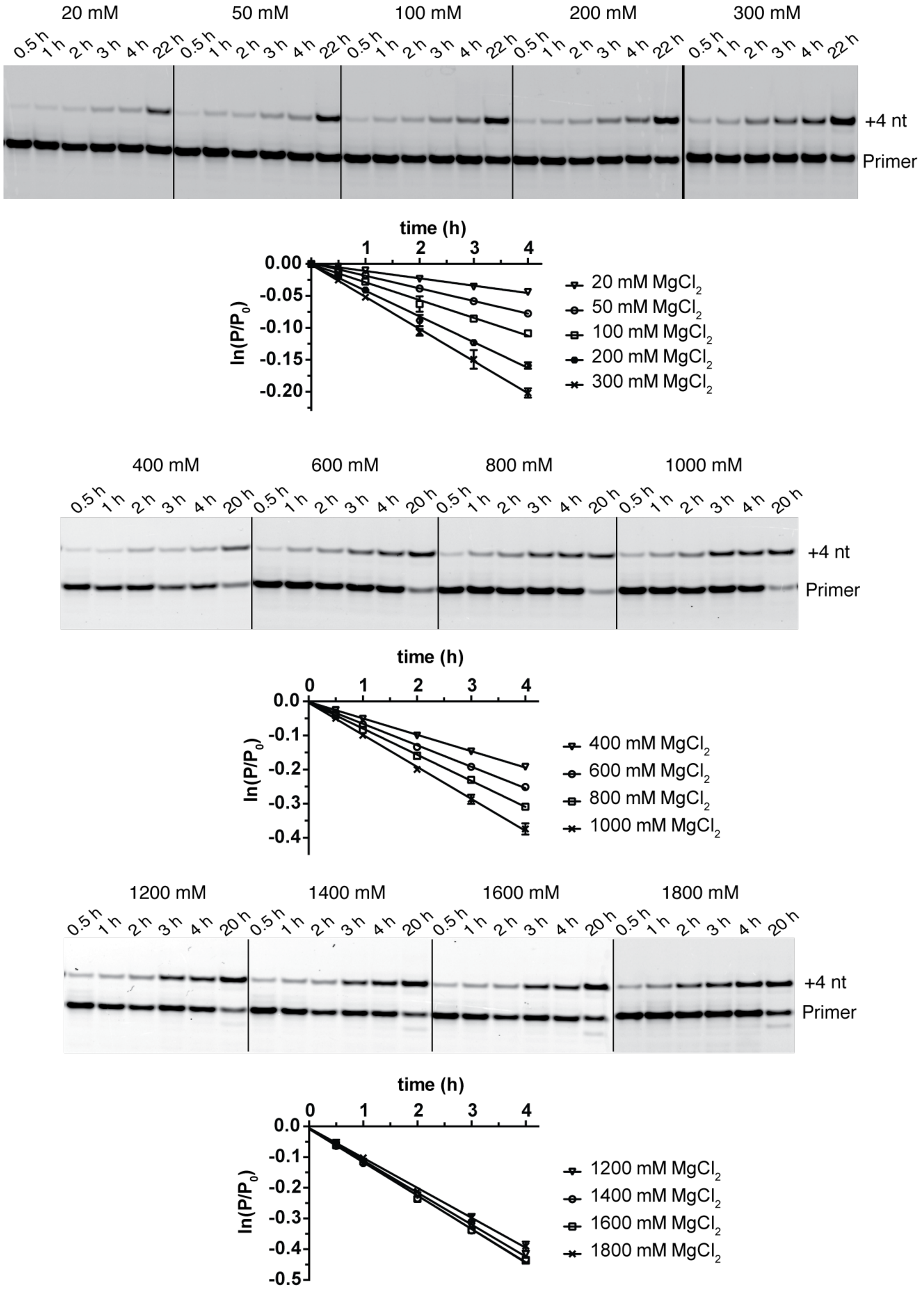


**Fig. S4**

Representative PAGE data and plots of $\ln(P/P_0)$ vs. time for the ligation kinetics described in Figure S3a.

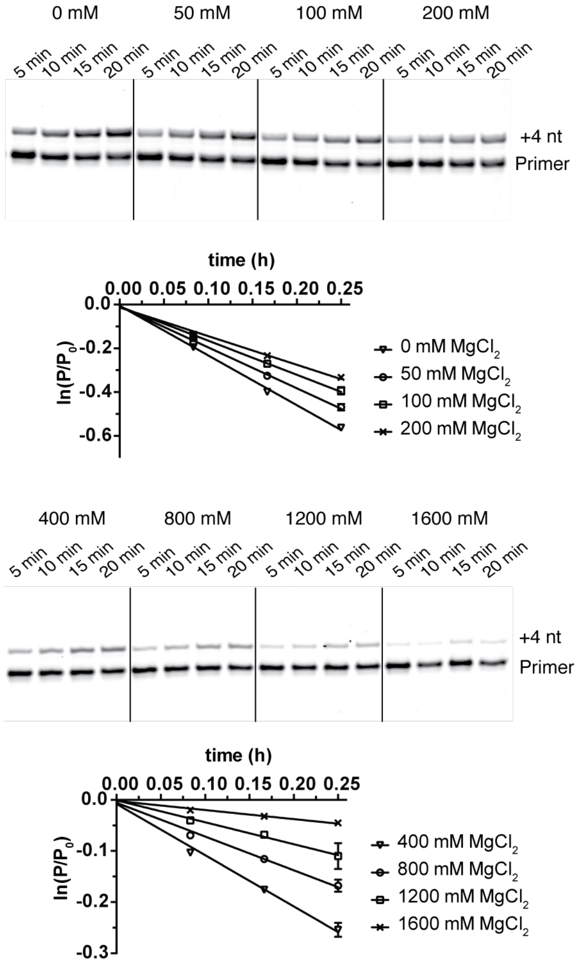


**Fig. S5**

Representative PAGE data and plots of $\ln(P/P_0)$ vs. time for the ligation kinetics described in Figure S3b.

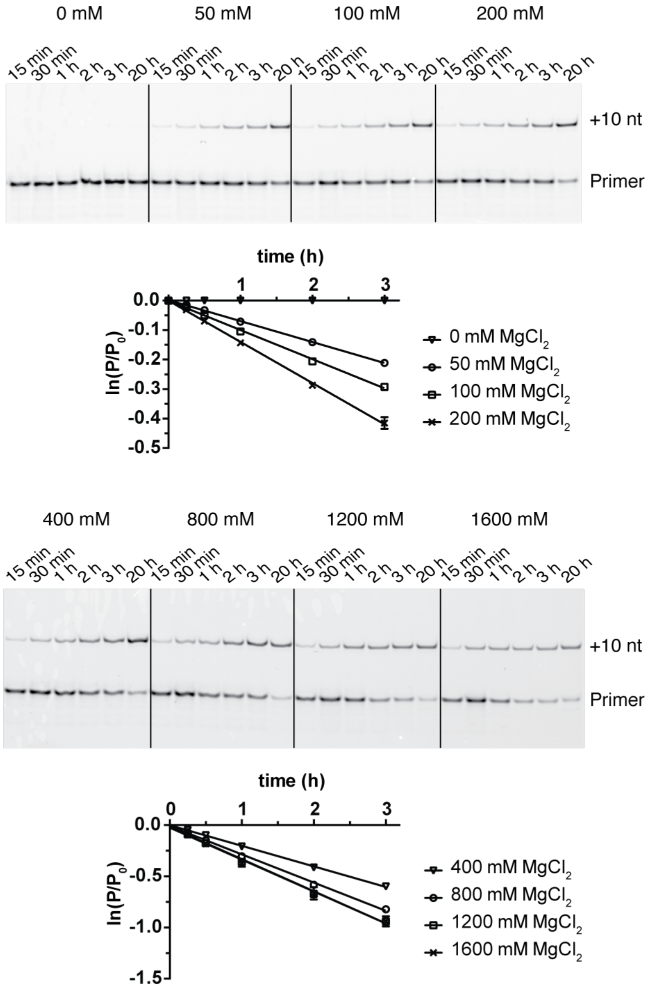


**Fig. S6**

Representative PAGE data and plots of $\ln(P/P_0)$ vs. time for the ligation kinetics described in Figure S3c.

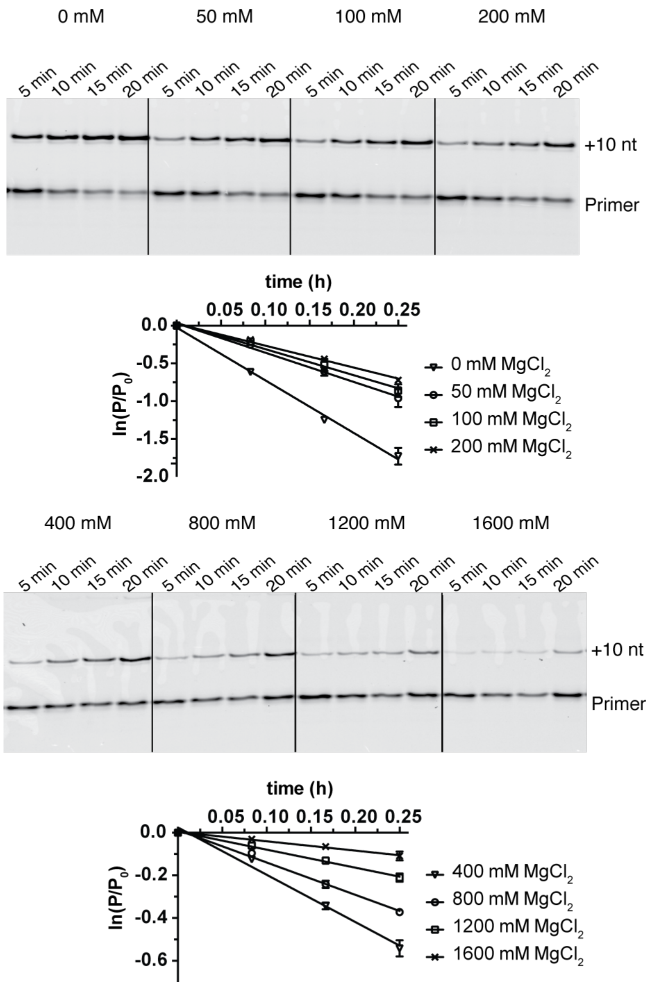


**Fig. S7**

Representative PAGE data and plots of $\ln(P/P_0)$ vs. time for the ligation kinetics described in Figure S3d.

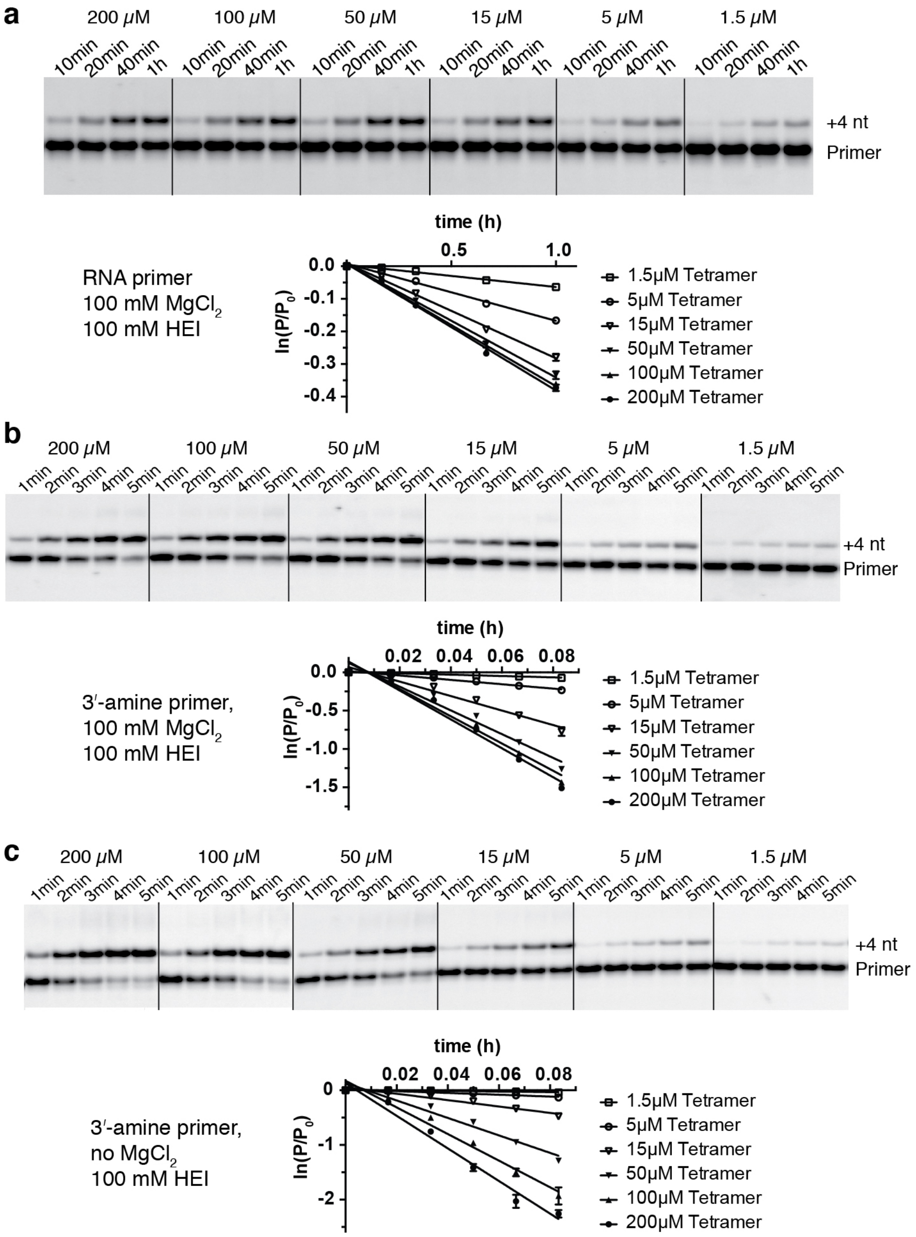


**Fig. S8**

Representative PAGE data and plots of $\ln(P/P_0)$ vs. time for the ligation kinetics described in Figures 1d and 1e.

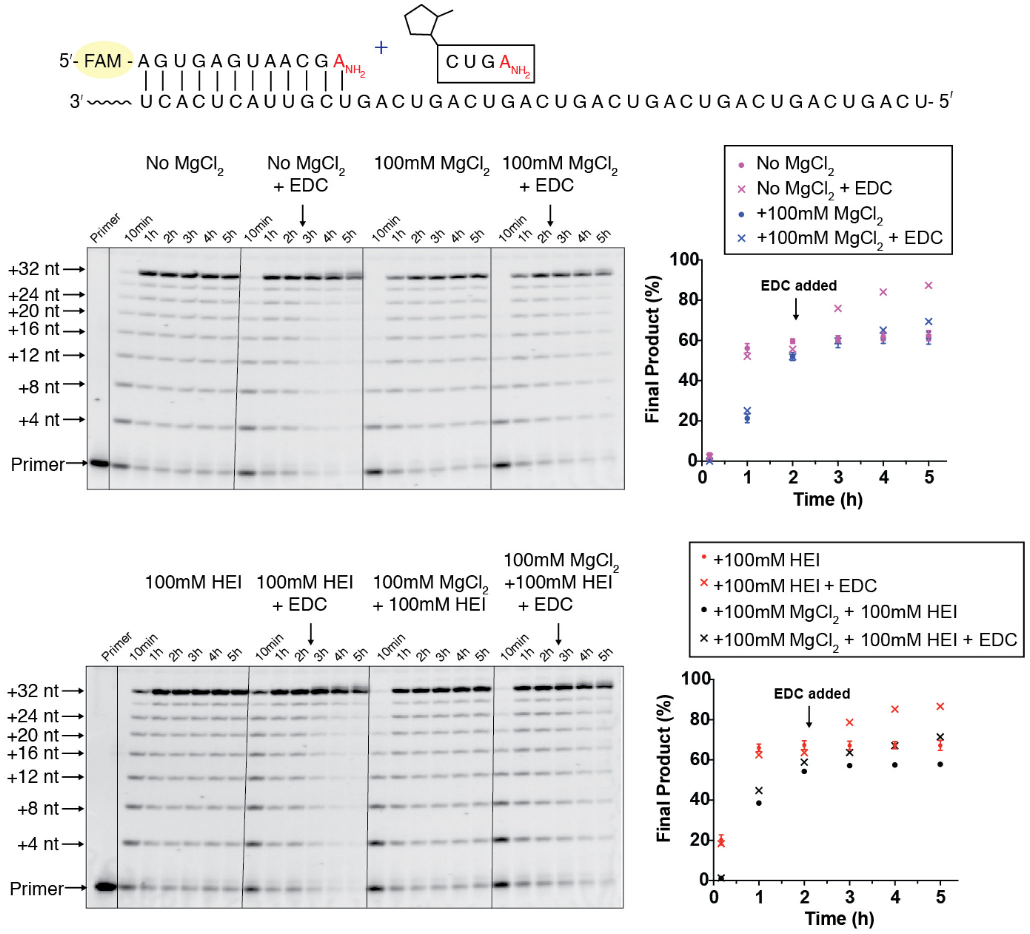


**Fig. S9:**

Ligation reactions as in Figure 2a except with the addition of 150 mM EDC after the 2h, 3h and 4h time points.

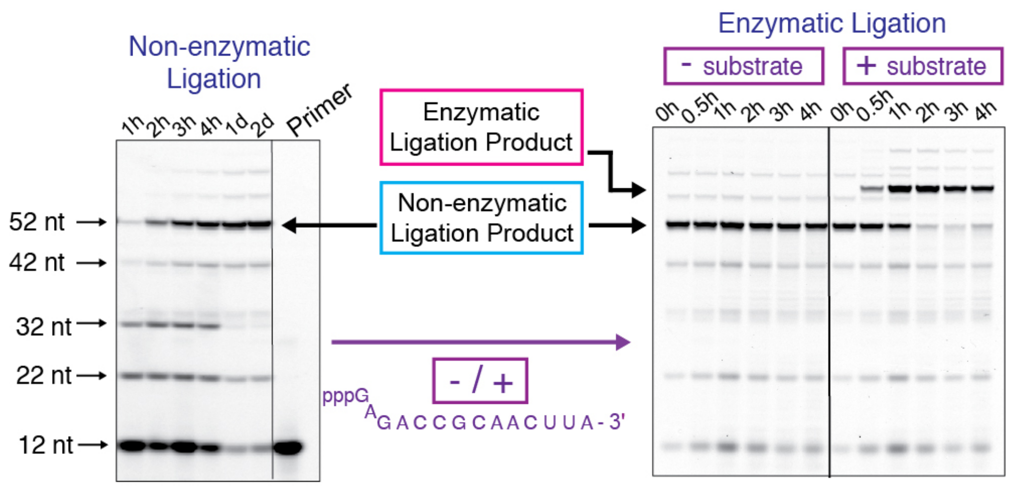


**Fig. S10:**

Assembly and activity of an RNA ligase ribozyme. Left PAGE: assembly of the RNA ligase *via* DNA splinted non-enzymatic ligation. The reaction contained 18 µM primer, 25 uM of each 2-methylimidazole activated oligomer as shown in Figure 4, 25 uM of 4 DNA splints (5′-AAGTGATAAC-3′, 5′-CTTACGTAAC-3′, 5′-CAAAGTGTTA-3′, 5′-TCAACCCATC-3′), 200 mM HEPES, pH 8.0, 100 mM NaCl, 1 mM EDTA, 50 mM $MgCl_2$ and 100 mM HEI. The ligation reaction was carried out on ice. Right PAGE: activity of the product ligase. The non-enzymatic ligation reaction mixture was diluted 10-fold into the ligase reaction buffer of 50 mM bis-tris propane, pH 8.5, 25 mM $MgCl_2$, with or without 3 µM of the illustrated 5′-triphosphate substrate oligonucleotide. The ligase reaction was performed at 48°C.

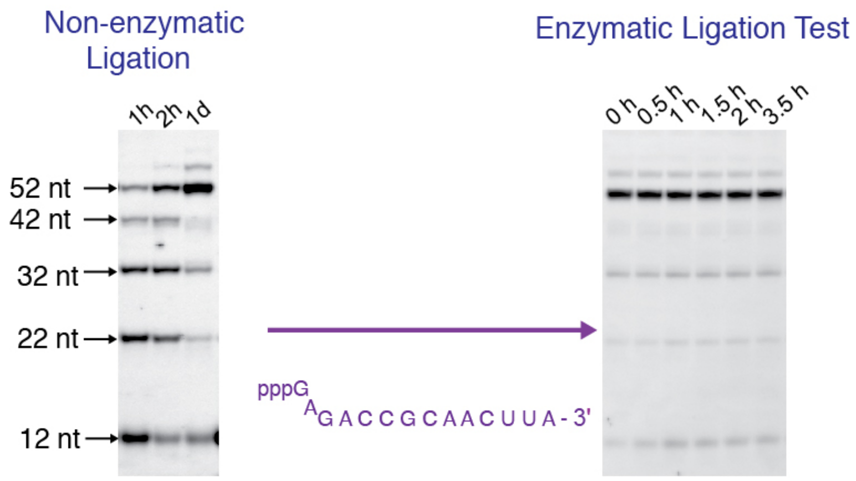


**Fig. S11:**

Assembly and lack of activity of an RNA ligase ribozyme in the presence of RNA splints. Left PAGE: efficient RNA splinted assembly of an RNA ligase ribozyme *via* non-enzymatic ligation. The reaction contained 20 μM primer, 25 uM of each 2-methylimidazole activated oligomer as shown in Figure 4, 25 uM of 4 different RNA splints (5′-AAGUGAUAAC-3′, 5′-CUUACGUAAC-3′, 5′-CAAAGUGUUA-3′, 5′-UCAACCCAUC-3′), 200 mM HEPES, pH 8.0, 100 mM NaCl, 1 mM EDTA, 50 mM $MgCl_2$ and 100 mM HEI. The ligation reaction was carried out on ice. Right PAGE: the product ligase is not functional. The non-enzymatic ligation reaction mixture was diluted 10-fold into the ligase reaction buffer of 50 mM bis-tris propane, pH 8.5, 25 mM $MgCl_2$, with 3 μM of the illustrated 5′-triphosphate substrate oligonucleotide. The ligase reaction was performed at 48°C.

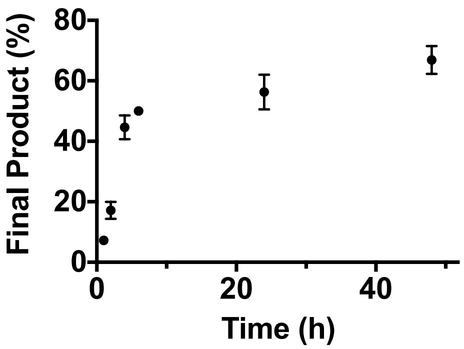


**Fig. S12:**
Yield of full-length RNA ligase assembly by non-enzymatic ligation using wobble mis-matched RNA splints, as shown in Figure 4.
